# Ethanolamine improves colonic barrier functions and inflammatory immunoreactions via shifting microbiome dysbiosis

**DOI:** 10.1101/2020.07.09.196592

**Authors:** Jian Zhou, Xia Xiong, Dan Wan, Hongnan Liu, Yirui Shao, Yuliang Wu, Xiali Huang, Chanfeng Peng, Pan Huang, Lijun Zou, Yulong Yin

## Abstract

Ethanolamine(EA) often occurs at a relatively high concentration within the inflamed gut of IBD patients. To investigate the role of EA in colonic inflammation and host-microbiome dysbiosis, thirty-six ICR mice were treated with 3% DSS for a week to generate acute intestinal inflammation and then supplied with 0μM, 500μM (LowEA), and 3000 μM (HighEA) in drinking water for two weeks, after that,16s RNA sequencing was applied in characterizing the changes in colonic microbiota driven by different EA levels. An inflamed colonic organoid model via 3% DSS treatment was also established for further verification of these in vivo findings.EA significantly reduced proximal colonic crypt depth but increased distal colonic villus height in HighEA group. The protein and mRNA expression of occludin and Reg3β, BD1, BD2, and MUC2were significantly up-regulated in EA treated groups. EA decreased mucosal inflammation-related cytokines levels (IL1, IL6, IL17, TNFα, and INFγ) and increased the significantly increased concentration of sIgA. Serum aspartate aminotransferase and alanine aminotransferase were significantly down-regulated in the highEA group. EA increased the relative abundance of *Blautia, Roseburia, Lactobacillus, Faecalibaculum, Candidatus_Saccharimonas, Alloprevotella*, and *Lachnoclostridum*.and thus microbial metabolic pathways including *Oxidative phosphorylation, Lipopolysaccharide biosynthesis, Arginine and proline metabolism, Folate biosynthesis*, and *Biotin metabolism* were more abundant in LowEA group than those in control. EA up-regulated the protein or mRNA expression of TLR4/MyD88 in colonic tissues and the DSS-treated colonic organoid model. This study firstly demonstrated that ethanolamine in altering host-microbiome dysbiosis, which may provide new insights into the role of dietary lipids in IBD.

**Importance:** Inflammatory bowel disease (IBD) affects ~3.1 million people in the USA and is increasing in incidence worldwide. IBD pathogenesis has been associated with gut microbiome dysbiosis characterized as a decrease in gut microbial diversity. Extensive works have demonstrated the roles of dietary fiber, short-chain fatty acids, and aromatic amino acids in altering the composition of gut microbiota to restore immune homeostasis and alleviate inflammation via diverse mechanisms in IBD. However, little is known about essential sphingolipids like ethanolamine (EA), an essential compound in the CDP-ethanolamine pathway for phosphatidylethanolamine (PE) in both intestinal cells and bacteria. PE synthesis deficiency can ultimately result in a loss of membrane integrity and metabolic disorders in IBD. Our results demonstrate that ethanolamine could improve colonic barrier functions and inflammatory immunoreactions via shifting microbiome dysbiosis, which provides new insights into the role of dietary lipids in IBD.

## 1. Introduction

Nutrient signals, microbiome dysbiosis, and host immunoreactions have a pivotal role in the development of inflammatory bowel diseases (IBD) and colorectal cancer^1–4^. The gastrointestinal tract not only acts as the primary site of food digestion and nutrition absorption but also provides an essential interface for the interactions of gut microbiota and immune systems^5^. Dramatic alterations in the composition of gut microbiota and intestinal barrier dysfunction have been found in IBD Patients^3, 6^. Nutrient signals including gut microbial-derived metabolites, aromatic amino acids, short-chain fatty acids and activate compounds can directly impact the establishment of the tumor-associated microenvironment and contribute to host-microbial interactions in the progress of the colorectal cancer^3–4^. Extensive works have demonstrated the roles of L-arginine^7^ and tryptophan^8^, and short-chain fatty acids like butyrate^9^ and acetate^10^ in altering host-microbial interactions to restore gut homeostasis and alleviate intestinal inflammation through diverse mechanisms^3^. However, little is known about essential sphingolipids like ethanolamine (EA), an essential compound in the CDP-ethanolamine pathway for phosphatidylethanolamine (PE) synthesis^11^. PE synthesis deficiency can ultimately result in a loss of membrane integrity and metabolic disorders in inflammatory diseases^10, 12^. The gastrointestinal tract keeps a physiological concentration of EA at 1~2mM via daily diet intake and internal recycling of PE to maintain intestinal metabolic homemstatsis^13^.

IBD patients consistently suffered from chronic inflammation and held more opportunities to develop cancer^14^. Lipid-mediated host-microbial interactions in metabolism and immune reactions are implicated in chronic inflammatory diseases like IBD^15^. Ethanolamine has been detected at a relatively high concentration within the inflamed gut of both IBD patients and rodent models as the hydrolysate of PE^16–17^. Ethanolamine can be utilized by both intestinal epithelial cells^11^ and bacteria like *Enterococcus faecalis*^18^ that hold the EA utilization (*eut*) genes via the CDP-EA pathway^10^, which confers it with the potential role in mediating the cross-talk between the intestinal epithelium and gut microbiota. A recent study highlighted that EA could promote the mesenchymal-to-epithelial transitions via a CDP-EA-Pebp1 dependent manner that ultimately leads to NF-κB inhibition^11^. Ethanolamine has been associated with the pathogenesis of *eut* pathogens such as *Salmonella*^16^ and enterohemorrhagic *Escherichia coli* (EHEC)^17^ that have been reported to promote colorectal carcinogenesis and tumor formation^2–3^. For instance, several studies have demonstrated that *S. Typhimurium* can sidestep nutritional competition with commensal bacteria by utilizing EA in inflamed gut^16, 19–20^. Toll-like receptors (TLRs) have a vital role in mucosal immune responses to gut bacteria, and the TLR4 expression was always dramatically up-regulated in the intestines of IBD patients^21^. Myeloiddifferentiationfactor (MyD) 88 holds its essential role in the regulation of innate gut immunity, and it is the direct downstream of TLRs and cytokine receptors^22^. Nutrient signals such as peptidoglycan and lipopolysaccharide can activate TLR4-MyD88 dependent or independent pathways to regulate the expression of antimicrobial proteins like the Reg3 protein family that ultimately reprogramme the gut microbiome in IBD^23–24^. However, The role of EA on TLR4/MyD88 signaling to restore microbiome dysbiosis under inflammatory conditions remains unknown.

Our previous study has preliminarily demonstrated that 500~1000 mM supplementation of EA could alleviate weaning stress via re-programing gut microbiota^13^. To investigate the potential role of Etn as a nutrient signal in microbial-host interactions, the impact of Etn on colonic morphology, antimicrobial protein expression, inflammation-related cytokines, serum indicators, and gut microbiome were investigated.

## 2. Materials and methods

### 2.1 Animals and Experimental Design

All experiments were approved by the Animal Care and Use Committee of the Institute of Subtropical Agriculture, Chinese Academy of Sciences^25^. Thirty-six male ICR mice (three-week-old) were obtained from the SLAC Laboratory Animal Central (Changsha, China). All mice were housed under standard conditions, in pathogen-free colonies (temperature, 22 ± 2 °C; relative humidity, 50 ± 5%; lighting cycle, 12 h/d), with free access to food and water. Intestinal colitis was induced in all mice by administration of 3% DSS (MP Biomedicals Shanghai, Co., Ltd. molecular weight=165.192 g/mol) in drinking water for seven days as previously described^26^. After that, thirty mice with colitis were obtained and randomly assigned into three groups (n =8) including the control group (Control) without EA (EA, Sigma-Aldrich,CAS141-43-5) supplements and two treatment groups that were supplemented with 500μM EA (LowEA) and 3000 μM EA(HighEA) in drinking water for two weeks (Figure 1A).

**Figure 1.**
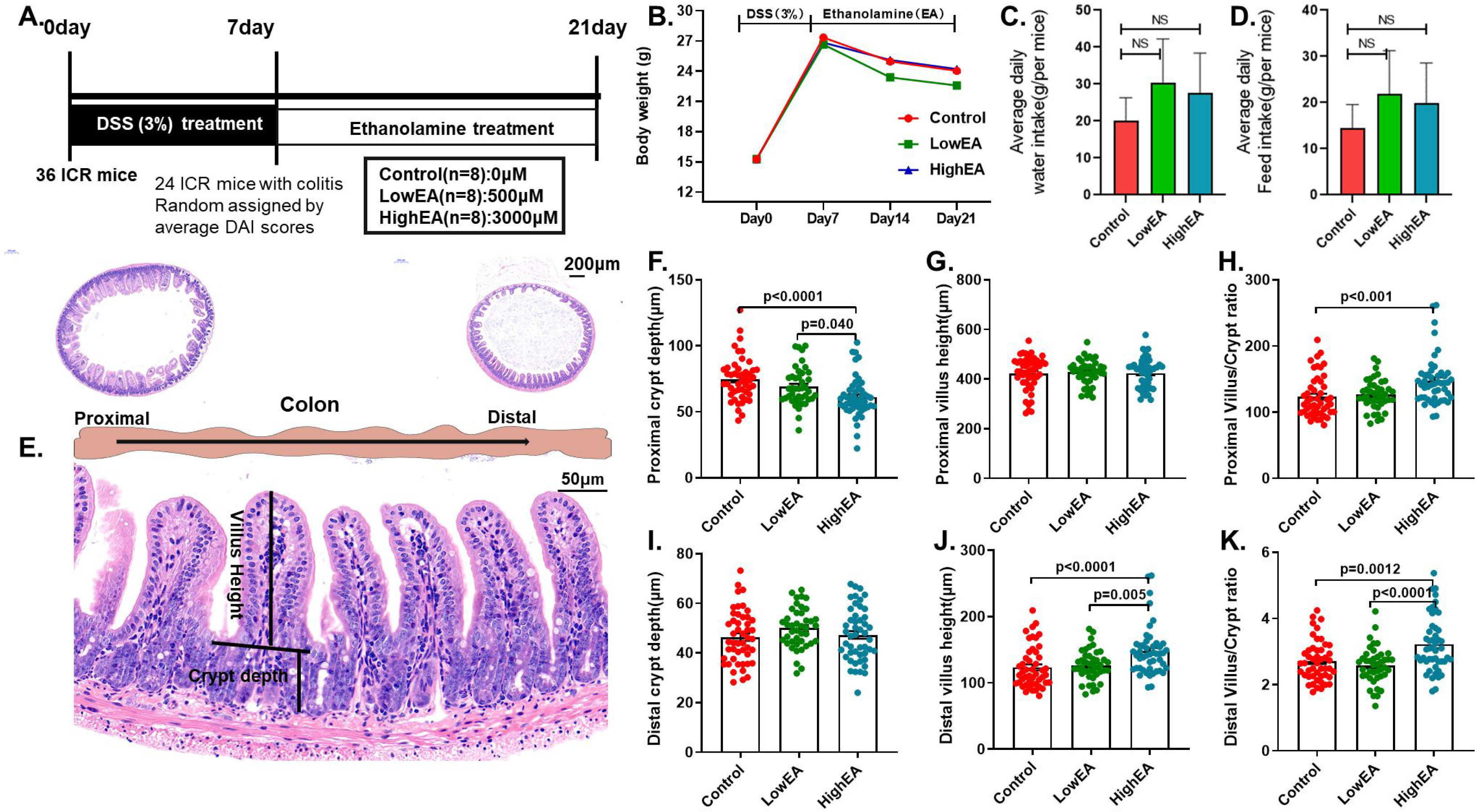
Ethanolamine treatments altered the colonic morphology of DSS-treated mice. **A.** DSS (3%) in drinking water was exploited in generating intestinal inflammation, and then different levels of ethanolamine were supplied in the same way. **B.** Ethanolamine treatment reduced the bodyweight of mice with colitis. **C-D.** Ethanolamine treatment did not affect average daily food and water intake. E. Both proximal and distal colon morphology of mice were investigated. **F-H**. Ethanolamine significantly reduced the crypt depth and crypt/villus ratio of the proximal colon. **I-K.** Ethanolamine increased the villus height and villus/crypt ratio of the distal colon.

### 2.2 Colonic crypt isolation and DSS-treated organoid culture

Mouse colonic crypts were harvested from three male 8-week-old ICR mice and cultured, as previously described^27^. In brief, 3 cm segments of the colon were minced and washed with cold DPBS(Stem cell,cat#37350) for 15~20 times until the supernatant is clear. The tissue pieces were digested in Gentle cell dissociation reagent (Stem cell,cat#07174) for 20~30min on a rocking platform at 20 rpm. The crypt compartment was collected by centrifugation and then washed twice with DMEM/F12 media(Stem cell,cat#36254). Then crypt pellets were resuspended with 25μl matrigel matrix (Corning,cat#356231) and 25μl medium per well and plated onto a 24-well tissue culture plated with complete mouse organoid growth medium(Stemcell, cat#06000) 10 μM Y-27632 ROCK inhibitor(Stem cell,cat#72302). After a week, mature colonic organoids were resuspended with matrigel matrix and medium as well as 3% DSS (MP Biomedicals,cat# 02160110) to form matrigel domes (Figure 8A) and cultured as previously reported methods^28^.

### 2.3 Morphological analysis

Cross-sections of tissue samples from each group were preserved in 4% formaldehyde, and glutaraldehyde mixing fixative was prepared using standard paraffin embedding techniques. Then samples were sectioned at 5μm thickness and were stained with hematoxylin and eosin as previously described^29^. Villus height (VH) and crypt depth (CD) were measured under a light microscope at 40× magnification using an image processing and analysis system (Leica Imaging Systems, Cambridge, UK). A minimum of 10 well-oriented, intact villi was measured from the crypt mouth to the villus tip and all measurements were made in 10 μm increments, and the count was repeated three times for each section per sample.

### 2.4 Immunohistochemistry Assay

Tissue samples from colons were cut into 4-μm sections and processed for immunohistochemical staining as described^30^. And then, these samples were incubated with a primary antibody anti-occludin(Abcam,cat#ab31721) overnight at 4°C and then with poly-horseradish peroxidase-conjugated occludin for 60min at 22±4°C. Subsequently, the avidin-biotin-peroxidase complex and the substrate 3,3’-diaminobenzidine were applied for 2min and the samples were analyzed.

### 2.5 Enzyme-Linked Immunosorbent Assay

Levels of IL1, IL6, TNFα,IL17,INFγ,sIgA,IL10, and IL22 in colonic segments were measured using ELISA kits according to the manufacturer’s instructions as previous studies^13^. The ELISA kits are from (Meimian, Yutong Biological Technology Co., Ltd, Jiangsu, China). Briefly, supplied diluent buffer in the kits was used to dilute standards and serum samples. Next, 100 μL of the sample or standard in duplicate was added to the wells of a microtitre plate precoated with wash antibody. Diluent buffer was applied as a negative control. The plates were incubated for 2 hours at 37°C. After incubation, 100 μL of biotin-antibody was added to each well after removing the liquid and incubated for 1 hour at 37°C. All wells were washed three times, with 200 μL volume of wash buffer. Next, 100 μL horseradish peroxidase-avidin was added to each well for 1 hour at 37°C. After a final wash, 90 μL of the supplied TMB substrate was added and incubated for thirty minutes in the dark at 37°C. The reaction was stopped with 50 μL of the supplied stop solution, and absorbance was measured at 450 nm with a spectrophotometer.

### 2.6 Serum biochemical analysis

Serum levels of total bile acid(TBA) were detected by the fully automatic biochemical analyzer(Shenzhen Mindray, BS-190), as described in our previous studies^25^. Serum aspartate aminotransferase (AST), alanine aminotransferase (ALT) and dual amine oxidase(DAO) were determined via commercial kits, and all determinations were done in triplicate and performed according to the manufacturer’s instructions^25^.

### 2.7 Quantifications Real-Time PCR

Total RNA of jejunal mucosa was extracted using TRIZOL reagent (Invitrogen, Carlsbad, CA). The first-strand cDNA was then synthesized using a reverse transcription kit (TaKaRa, Dalian, China) following the manufacturer’s instruction, as described in our previous study^29^. qPCR was performed using primers shown in the supplemental materials (Sup_Table 1). Briefly, 1 μL cDNA template was added to a total volume of 10 μL assay solution containing 5 μL SYBR Green mix,0.2 μL Rox, 3 μL deionized H2O, and 0.4 μL of each of the forward and reverse primers. The comparative Ct value method was used to quantitate mRNA expression relative to β-actin. All samples were run in triplicate and the average values were calculated.

### 2.8 Western Blot Analysis

Total proteins from intestinal tissues were extracted with a commercially available kit using lysis buffer containing 1‰ DTT, 5‰ PMSF and 1‰ protease inhibitor (KeyGEN BioTech, Nanjing, China). Following intermittent vortexing for 20 min on ice, samples were centrifuged at 13,000 rpm for 15 min at 4 °C. Protein content was determined using the BCA assay (Pierce Biotechnology, Rockford, IL, USA), and Western blotting analysis was performed. Briefly, 20 μg protein per lane was separated by SDS-PAGE and blotted onto nitrocellulose membranes. Primary antibody against occluding (Abcam,cat#ab31721),anti-TLR4(Abcam,cat#13556) and anti-MyD88(Abcam,cat#133739) and recombinant anti-beta actin antibody (Abcam,cat#ab115777) were incubated with the membrane overnight at 4 °C. After incubating overnight with the secondary antibody, the membrane was exposed to EZ-ECL (Biological Industries, Cromwell, CT, USA) for protein band detection, and all original images of WB results could be found in supplemental materials (Sup_Figure 1).

### 2.9 Microbial sequencing of colon contents

Total bacterial DNA from approximately 0.25 g of colon contents was extracted using a QIAamp DNA Stool Mini Kit (Qiagen, Hilden, Germany) according to the manufacturer’s instructions as described^13^. The diversity and composition of the bacterial community were determined by high-throughput sequencing of the microbial 16S rRNA genes. The V4 hypervariable region of the 16S rRNA genes was PCR amplified using 515F: 5’-GTGCCAGCMGCCGCGGTAA-3’ and 806R: 5’-GGACTACHVGGGTWTCTAAT-3’ primers, Illumina adaptors, and molecular barcodes. Paired-end sequencing was performed on the Illumina HiSeq 2500 platform (Novogene, Beijing, China) to obtain raw 16s data. The assembled HiSeq sequences obtained by this research were submitted to the NCBI’s Sequence Read Archive with project ID: PRJNA551369 for open access.

### 2.10 Bioinformatic analysis and metagenomic predictions

Raw 16S data sequences The first-strand before being screened and assembled using the QIIME (v1.9.1) and FLASH software packages as previously described^25^. UPARSE (v7.0.1001) was used to analyze high-quality sequences and determine OTUs. Subsequently, high-quality sequences were aligned against the SILVA reference database (https://www.arb-silva.de/) and clustered into OTUs at a 97% similarity level using the UCLUST algorithm (https://drive5.com/usearch/manual/uclust_algo.html). Each OTU was assigned to a taxonomic level with the Ribosomal Database Project Classifier program v2.20 (https://rdp.cme.msu.edu/), as previously reported^13^. Functional metagenomes of all samples were predicted using Tax4Fun R packages(http://tax4fun.gobics.de)^31^. Further statistical interrogation and graphical depictions of microbiome data were performed by R software (v3.6.0) and related packages.

### 2.11 Statistical Analysis

Data were analyzed using the General Linear Model (GLM) procedure in SAS (SAS Institute Inc., Cary, NC) to identify significant treatment effects and interactions. All data were presented as Least Squares means plus pooled SEM. The Tukey multiple comparison tests were used to evaluate the differences among the treatments. Probability values ≤ 0.05 were taken to indicate statistical significance.

## 3. Results

### 3.1 Impacts of ethanolamine treatment on body weight and colonic morphology

There were no differences in the BW, ADF, and water intake among the three groups (**Figure 1 C-D**). Compared with control group, Etn decreased the proximal colonic crypt depth (*P* < 0.0001, Figure 1 F-H) and increased villus/crypt ratio (*P* < 0.0001). Meanwhile, EA significantly increased the villus height and villus/crypt ratios of distal colon (*P* < 0.05).

### 3.2 Ethanolamine altered the composition of colonic microbiota

Microbial diversity indicators (Shannon and Simpson)were significantly in the low EA treated group than the control group (*P* < 0.0001; Figure 2 A&B). Besides, microbial abundance represented by Chao1 (*P* = 0.014, Figure 2 C) and ACE (Figure 2 D) were significantly high in the low EA treated group than the high EA treated group. Phylogenetic diversity indicator PD_whole_tree was remarkably low than both the control group and the low EA treated group (Figure 2 E). But no difference in observed species was found (Figure 2 F).

**Figure 2.**
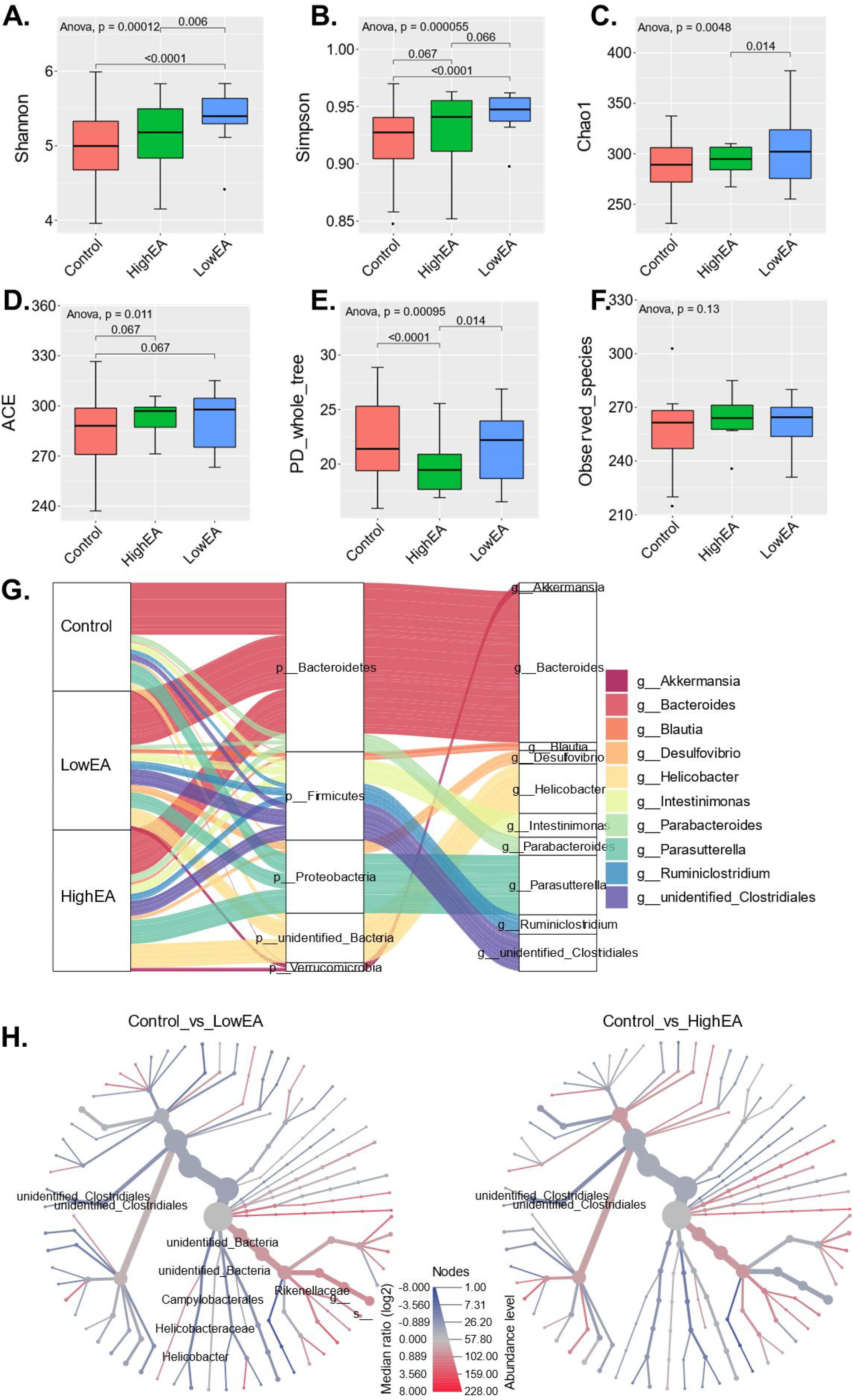
Impacts of EA treatments on the colonic microbiome. **A-B**. Shannon and Simpson indicators for microbial diversity. **C-D.** Chao1 and ACE indicators for microbial abundance. **E.** The metric PD_whole_tree for phylogenetic diversity. **F.** Observed species in the microbiome. **G.** Top-ten most abundant phylum and genus of microbiota in both the control group and EA treated groups. **H-I.** Significantly altered bacteria driven by different EA levels. The heat tree analysis was applied to leverage the hierarchical structure of taxonomic classifications to quantitatively (using the median abundance) and statistically (using the non-parametric Wilcoxon Rank Sum test) depict taxonomic differences between microbial communities.

The dominant phylum constituted by *Bacteroides*, *Firmicutes*, *Proteobacteria*, and *Verrucomicrobia*, likewise, *Akkermansia, Bacteroides, Blautia, Desulfovibrio, Helicobacter*, and *Intesinimonas* are the most abundant at the genus level (Figure 2 G). Besides, the heat tree analysis leveraged the hierarchical structure of taxonomic differences between the control group and EA treated group. *Unidentified_Clostridiales* and *unidentified_bacteria* significantly altered in two branches as a comparison between the control group and the EA treated group (Figure 2 H). Meanwhile, only *unidenfied_clostridiales* significantly altered when compared to the control group and the high EA treated group (Figure 2 I).

Furthermore, heatmap analysis on the top 40 most-abundant genus was performed (Figure 3 A). Notably, *Blautia, Roseburia, Lactobacillus* hold more abundance in the high EA treated group while *Faecalibaculum, Candidatus_Saccharimonas, Alloprevotella*, and *Lachnoclostridum.et.al* showed to be more abundant in the low EA treated group. Lefse analysis of the whole microbiome showed that differences among groups mainly occurred at the downstream of the family level (Figure 3 B). In that case, the T-test analysis of microbial composition was also performed (Figure 3 C), and results presented that *Helicobateraceae, unidentified_Clostridiales* increased while *Rikenellaceae* were reduced in the low EA treated group (*P* < 0.05). Only unidentified_Clostridiales were significantly increased in high EA treated groups.

**Figure 3.**
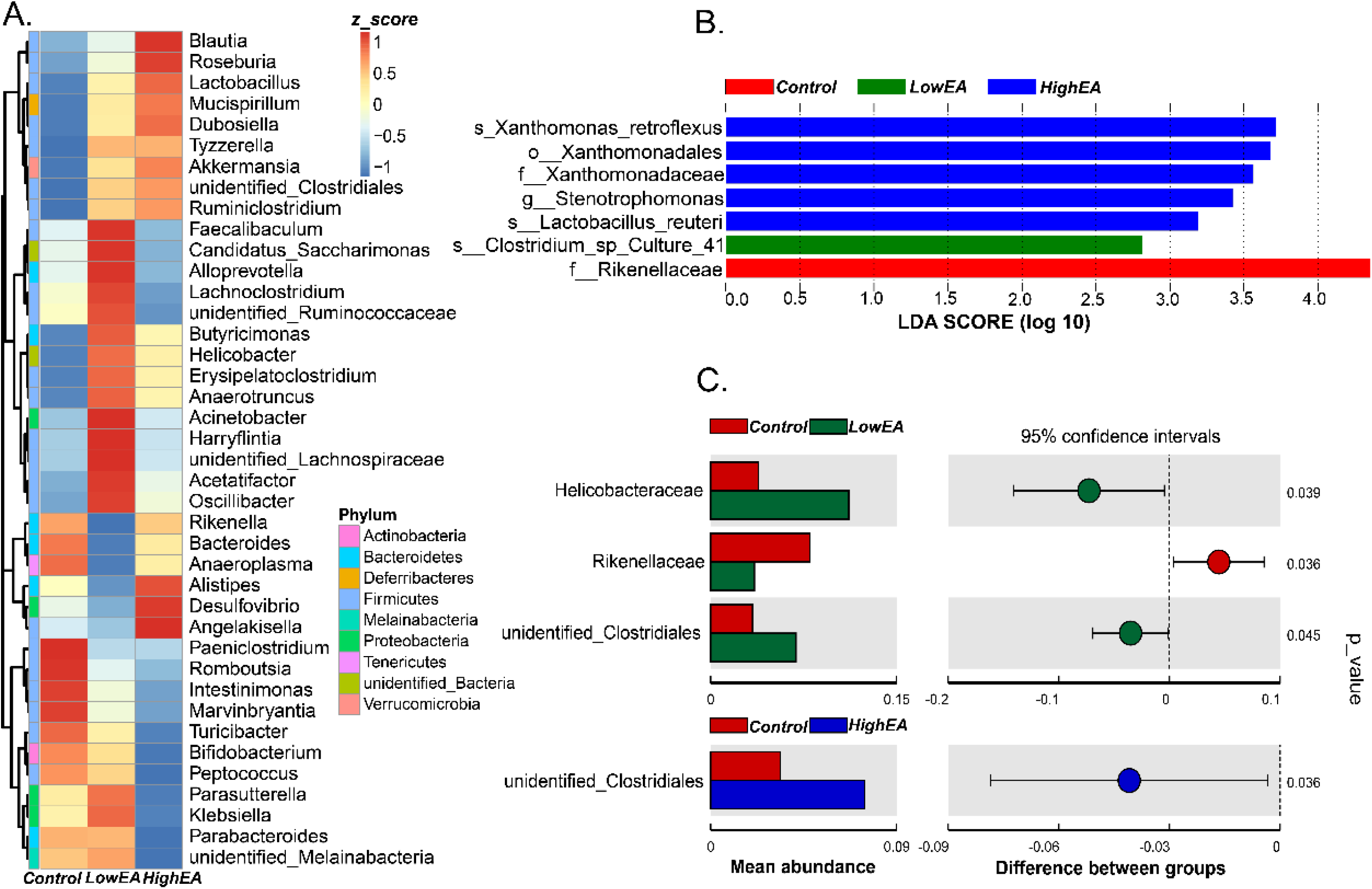
Microbial differences driven by different EA levels in the colon of DSS-treated mice. **A.** Genus composition of both the control group and EA treated groups. **B.** Lefse analysis of whole OTU in all groups (LDA score=2.0). **C.** Differences in microbial composition between control and EA treated groups at the genus level.

### 3.3 Metabolic functions driven by different EA levels

PCA analysis based on metagenome showed no apparent separations among groups (Figure 4 A). *Carbohydrate metabolism, Membrane transport, Replication and repair, Translation, Amino acid metabolism, Energy metabolism, Nucleotide metabolism, Glycan biosynthesis and metabolism, Metabolism of cofactors and vitamins* and *Signal transduction* were the top ten most abundant pathways at KEGG level 2 (Figure 4 B). Heatmap illustrated the composition of KEGG level 3 pathways that enriched in each group (Figure 4 C). *Glycan biosynthesis, Metabolism of cofactors and vitamins*, and *Nucleotide metabolism* showed more richness in the high EA group. However, no statistically significant differences between the control group and high EA treatment were observed at KEGG level 3. But significant variations between the control group and the low EA treated group were identified and that *Oxidative phosphorylation, Lipopolysaccharide biosynthesis, Arginine and proline metabolism, Folate biosynthesis* and *Biotin metabolism* were more abundant in the low EA group (Figure 4 D).

**Figure 4.**
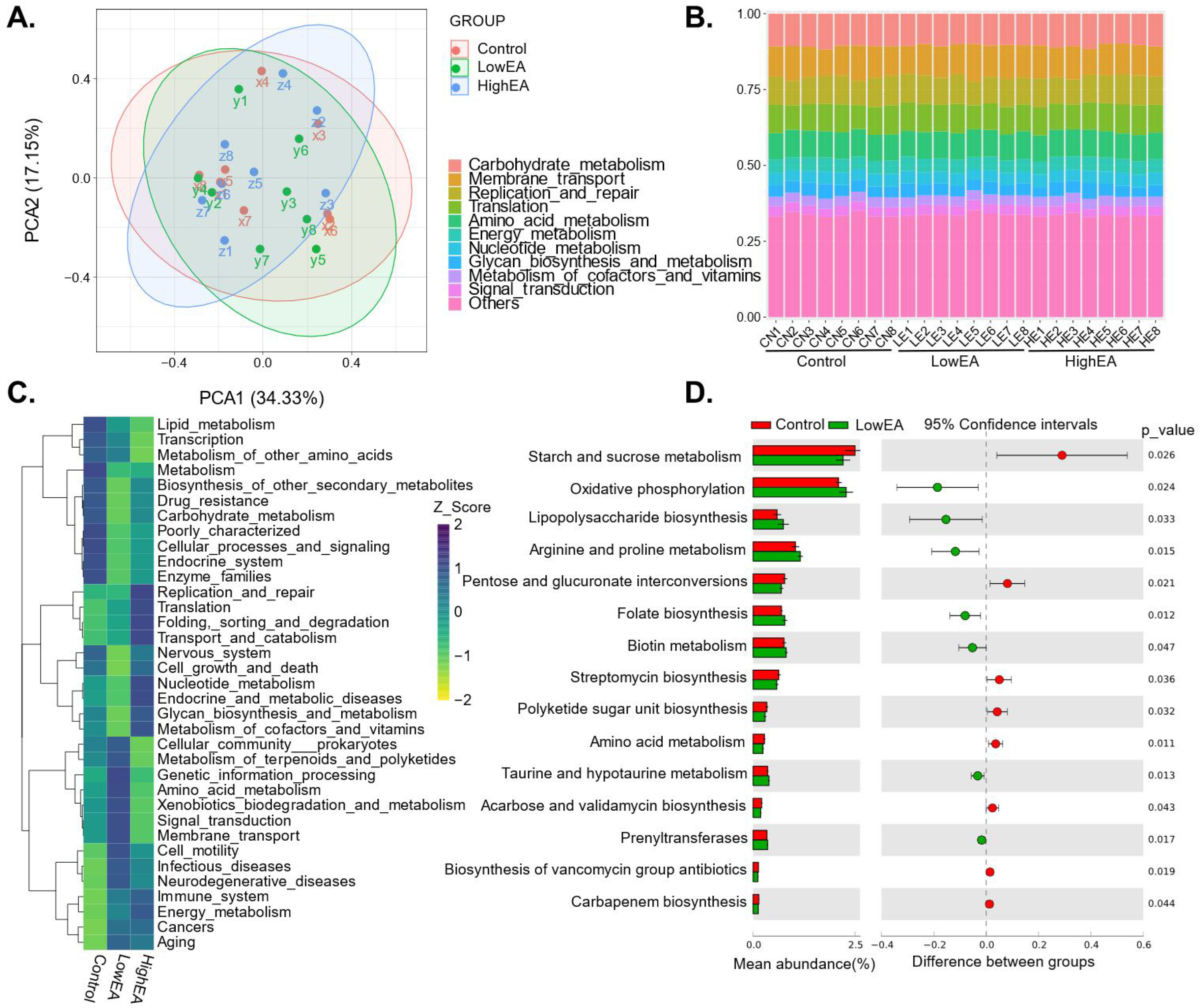
Metabolic functions of colonic microbiota driven by different EA levels in the DSS-treated mice model. **A.** PCA analysis of predicted KEGG pathways. **B.** Top-10 most abundant pathways at KEGG level-2. **C.** Composition of metabolic pathways of colonic microbiota at KEGG level-3. **D.** Differences in metabolic functions between control and EA treated groups at the KEGG level-3. Moreover, no statistically significant differences existed between the control group and high EA treatment

### 3.4 Impacts of EA treatment on body weight and colonic morphology

EA treatments had no impacts on body weight, average daily food and water intake (**Figure 5**). Compared with the control group, the proximal colonic crypt depth of both EA treated groups showed significant reductions (*P* < 0.0001, **Figure 5 F-H**)) and villus/crypt ratio significantly increased in the high EA treated group (*P* < 0.0001). Meanwhile, EA treatments significantly increased the villus height and villus/crypt ratios of distal colon (*P* < 0.05).

**Figure 5.**
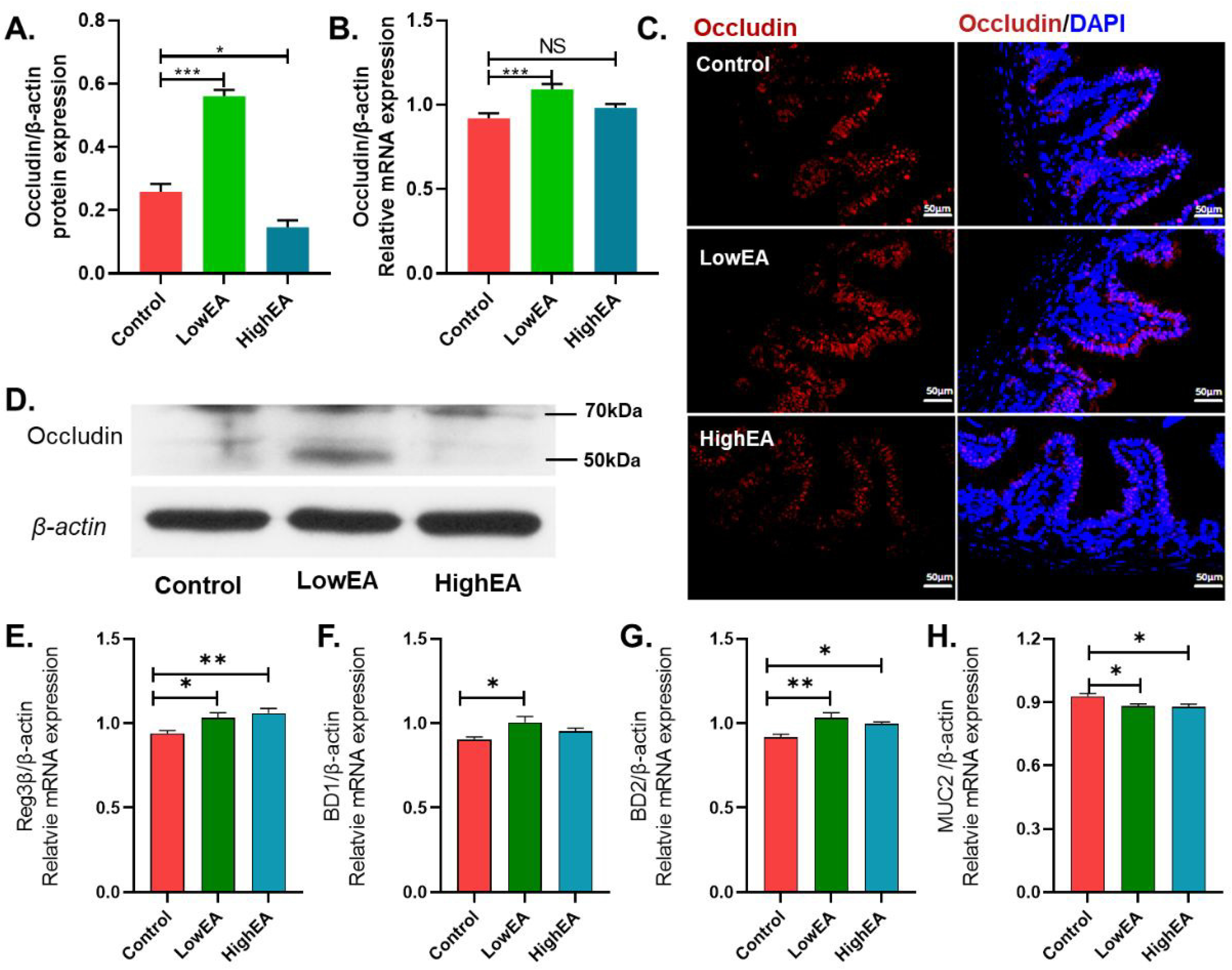
EA treatments altered the colonic morphology of DSS-treated mice. **A.** DSS(3%) in drinking water was exploited in generating intestinal inflammation, and then different levels of EA were supplied in the same way. **B.** EA treatment reduced the bodyweight of mice with colitis.**C-D.** EA treatment did not affect average daily food and water intake. E. Both proximal and distal colon morphology of mice were investigated. **F-H**. EA significantly reduced the crypt depth and crypt/villus ratio of the proximal colon. **I-K.** EA increased the villus height and villus/crypt ratio of the distal colon.

### 3.5 Ethanolamine altered intestinal permeability and antimicrobial protein mRNA expression

The mRNA and protein expression of occludin in the low EA treated group were increased compared with those in control groups (*P* < 0.001, Figure 6 A-D). Antimicrobial protein Reg3β, BD2, and MUC2 mRNA were significantly up-regulated in EA treated groups(*P* < 0.005). Relative expression of BD1 was remarkably more significant in the low EA treated group than in the control group(*P* < 0.05, **Figure 6 F**).

**Figure 6.**
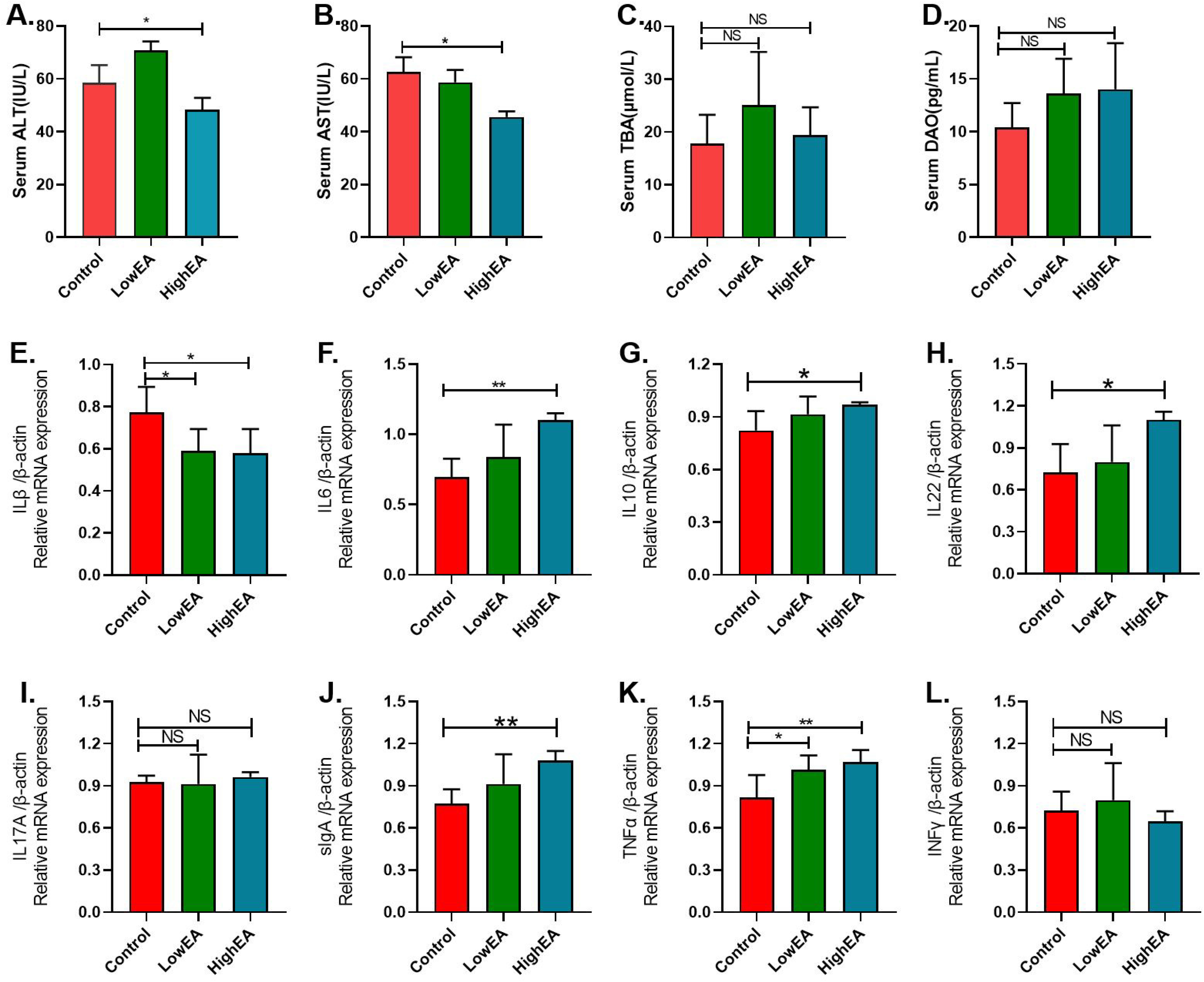
Impacts of EA on mucosal permeability and antimicrobial protein mRNA expression. **A-D.** Protein and mRNA expression of occludin.E-H,mRNA expression of Reg3β, BD1, BD2and MUC2.

### 3.6 Ethanolamine changed the production and mRNA expression of inflammation-related cytokines and Serum indicators

For mucosal inflammation-related cytokines in the colon, the concentration of IL1, IL6, IL17, TNFα, and INFγ showed a significant reduction in EA treated groups (*P* < 0.01). Conversely, sIgA was significantly up-regulated in both low and high EA treated groups (Table 1).

**Table 1.** Expression of inflammation-related cytokines driven by different EA levels

| Cytokines | Control | LowEA | HighEA | P-value |
| --- | --- | --- | --- | --- |
| pg/mg·pro | 0μM | 500μM | 3000μM |  |
| IL1 | 202.8±11.42<sup>a</sup> | 136.75±9.048<sup>b</sup> | 87.903±6.480<sup>c</sup> | <0.001 |
| IL6 | 150.8±8.595<sup>a</sup> | 110.9±11.29<sup>b</sup> | 71.59±4.911<sup>c</sup> | <0.001 |
| TNFα | 1185±33.09<sup>a</sup> | 889.4±72.60<sup>b</sup> | 566.2±30.21<sup>c</sup> | <0.001 |
| IL17 | 53.87±2.941<sup>a</sup> | 48.30±2.184<sup>a</sup> | 40.78±2.533<sup>b</sup> | 0.003 |
| INFγ | 115.1±6.708<sup>a</sup> | 85.15±6.030<sup>b</sup> | 40.22±3.054<sup>c</sup> | <0.001 |
| sIgA | 36.11±2.209<sup>b</sup> | 39.06±2.605<sup>ab</sup> | 45.18±1.935<sup>a</sup> | 0.022 |
| IL10 | 575.4±15.93 | 635.0±37.34 | 602.0±30.64 | 0.364 |
| IL22 | 54.23±1.963 | 52.88±2.895 | 52.66±3.367 | 0.913 |

**a,b** Values within a row with different superscript letters differ significantly at $p < 0.05$ .

The concentration of serum ALT and AST were significantly down-regulated in the high EA treated group compared with the control group (*P* < 0.05, Figure 7 A&B) and no differences in TBA and DAO concentration were found in this study (Figure 7 C&D). The mRNA expression of IL1β showed a significant reduction in the EA treated groups, and the expression of IL6, IL10, IL22, sIgA and TNFα mRNA were significantly up-regulated (*P* < 0.05, E-L).

**Figure 7.**
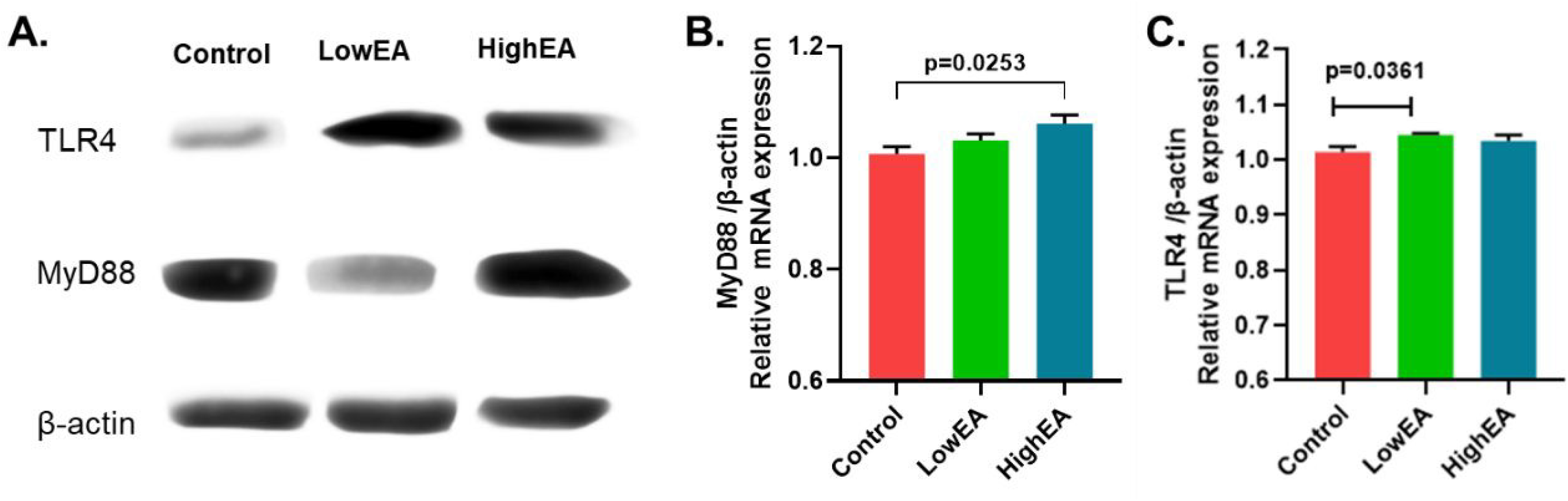
Impacts of EA on serum indicators and mRNA expression of cytokine-related genes. **A-D.** Impacts of EA on serum ALT, AST, TBA, and DAO. **E-L**. Relative mRNA expression of IL1β,IL6,IL10,IL22,IL17A,sIgA,TNFα and INFγ genes.

### 3.7 Ethanolamine promoted TLR4/MyD88 dependent signaling in inflamed colon tissues

As the dramatic changes in microbiota discussed above, we further determine whether EA altered gut microbiota composition rely on TLR4/MyD88 dependent signaling. Compared with control group, the protein expression of TLR4 in LowEA and MyD88 in HighEA were significantly up-regulated (Figure 8 A). The mRNA expression of MyD88 was up-regulated in the high EA treated group (Figure 8 A). However, while TLR4 only showed a significant increase in the low EA treated group (*P* < 0.05, Figure 8 C).

**Figure 8.**
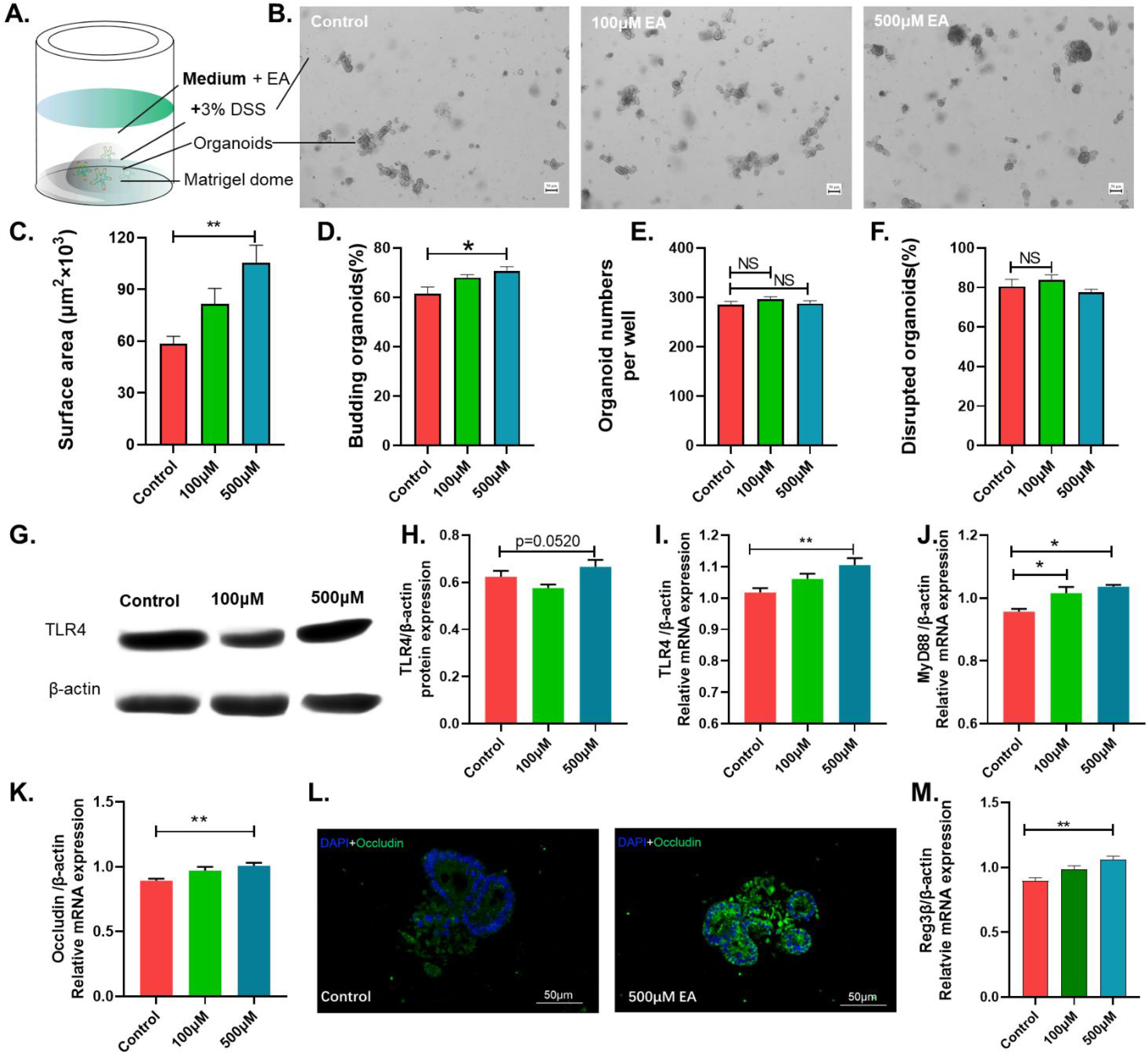
Impacts of EA on protein and mRNA expression of TLR4 and MyD88.

### 3.8 Ethanolamine increased the expression of TLR4 and occludin in the DSS-treated organoid model

To further determine the effects of EA on TLR4 expression and intestinal permeability, colonic organoids from intestinal crypts were directly derived from three male 8-week-old ICR mice for the establishment of DSS-treated organoid model (Figure 9 A&B). The preliminary experiment showed that organoids could not survive in a 3000μM EA environment. In that case, ethanolamine treatment with 0μM, 100μM, and 500μM were performed in the DSS-treated organoid model. 500μM ethanolamine supplementation in the organoid growth medium significantly increased the surface area and budding rate of colonic organoids (*P* < 0.05, Figure 8 C&D). TLR4 protein expression tended to be more significant in the 500μM EA treated group (Figure 9 G&H). The mRNA expression of TLR4 in 500μM ethanolamine treated group and MyD88 in both 100μM and 500μM ethanolamine treated group showed a significant increase (Figure 9 I-J). For mucosal permeability, occludin mRNA was significantly up-regulated in 500μM ethanolamine treated group (Figure 9 K-L). The relative mRNA expression of all antimicrobial proteins, including Reg3β, BD1, BD2, and MUC2, were also detected on colonic organoids, and only Reg3β showed a significant up-regulation in 500μM ethanolamine treated group (Figure 9 M).

**Figure 9.** Impacts of EA on TLR-4 and Occludin expression on DSS-treat colonic organoids. **A-B.** Establishing of DSS(3%) treated the colonic organoid model **C-D**. Impacts of EA treatments on the surface area and budding efficiency of colonic organoids. **E-F**. Organoid numbers per well and DSS disrupted organoid ratio. **G-H**. TLR4 protein expression in DSS-treated organoids driven by different EA levels. **I-J.** Relative mRNA expression of TLR4 and MyD88 genes. **K.** Occludin mRNA expression. **L.** Occludin imaging of the control group and 500μM group. **M.** Relative mRNA expression of antimicrobial proteins, only Reg3β significantly increased in 500μM group.

## Discussion

Nutrition signal to the host-microbiome interactions ultimately affect the pathological process and health outcomes^3–5, 15^. Dramatic alterations in gut microbiota and barrier dysfunction reflecting a different ecological microenvironment in IBD Patients and mice models compared with health conditions have been demonstrated by extensive works^1, 3–4, 6, 23^. Patients with Crohn’s disease showed reduced diversity of fecal microbiota and held the gut microbiome that dominantly constituted by *Proteobacteria, Actinobacteria, Firmicutes*, and *Bacteroidetes*^32^. Dietary intervention with optimal dietary components such as fiber, short-chain fatty acids, and tryptophan has been proved to be able to reduce colonic inflammation and cell proliferation and shift the progression of IBD through the gut microbiota^3, 33^. Sphingolipids like PE and EA have been detected as the most differentially abundant metabolite in stool and intestinal tissue resections from IBD patients^3, 15^. Here, we tried to demonstrate the role of ethanolamine as an essential lipid signal in mediating host-microbial interactions and its impacts on colonic barrier functions and inflammation.

The present study showed that 500 μM EA increased microbial diversity. However, higher concentration of EA (3000μM) decreased microbial abundance. Gut microbiome dysbiosis in IBD has been identified as a decrease in microbial diversity and abundance due to a shift in the balance between commensal and potentially pathogenic bacteria^34^. These changes indicated that EA could change intestinal microbial compostion of the DSS-treated mice. The dominant phylum in EA treated groups constituted by *Bacteroides, Firmicutes, Proteobacteria, and Verrucomicrobia, Akkermansia, Bacteroides, Blautia, Desulfovibrio, Helicobacter*. A recent study highlighted the role of *Bacteroides*-derived sphingolipids in host-microbe symbiosis and inflammation, where IBD patients have decreased sphingolipid production by *Bacteroides*, but increased host-produced sphingolipid abundance in gut^15^. *Akkermansia, Bacteroides*, and *Blautia* are associated with the production of beneficial microbial-derived metabolites such as short-chain fatty acids that have been proved to be a key regulator in IBD^3^. Studies have shown that short-chain fatty acids can inhibit histone deacetylases in colonocytes and intestinal immune cells to downregulate proinflammatory cytokines and induce apoptosis in cancer cell lines^1–2^. Ethanolamine can generate acetate^10^, that may be responsible for the beneficial role of EA in altering the gut microbiota. Here, we further analyzed the metabolic functions of the gut microbiome via applying theTax4fun algorithm package^31^. Significant variations between the control group and the low EA treated group were identified, and *Oxidative phosphorylation, Lipopolysaccharide biosynthesis, Arginine and proline metabolism, Folate biosynthesis* and *Biotin metabolism* were more abundant in the low EA treated group. These metabolic changes are coincident with studies that reported beneficial impacts of dietary fiber intervention in IBD^3, 34^. These results may indicate that EA could shift the composition and metabolic functions of colonic microbiota to restore microbial dysbiosis.

In this study, EA decreased the proximal colon crypt depth and increased the distal villus height. Previous study showed that Gut microbiota, including *Escherichia, Enterococcus, Bacteroides*, and *Clostridium genera*, could promote colorectal carcinogenesis by increasing aberrant crypt foci induced by 1, 2-dimethylhydrazine^3^. Under health conditions, the mucus layer covers the whole epithelial surface, which confers the lubrication, hydration, and protection of the underlying epithelial cells and the epithelial barrier integrity of intestines^36–37^. Tight junction proteins like occludin and ZO1 also contribute to the regulation of intestinal permeability that physically against the invasion of gut bacteria^37^. Antimicrobial proteins like BD1, BD2, and Reg3 family are often secreted by Paneth cells to clear up the pathogens and promote epithelial regeneration following intestinal infection^2, 23, 25^. However, IBD patients often suffered from prolonged enteric inflammation that ultimately damages intestinal epithelia, and subsequently leads to increased intestinal permeability that causing pathogenic infections^35^. For instance, *Akkermansia* recently has been proved to be a probiotic that may be beneficial to the IBD therapy, and direct administration of *Akkermansia* has been demonstrated to alleviate obesity-related metabolic disturbances and increase Muc2 production in DIO-mice, thus improving mucus layer thickness and intestinal permeability^38–39^. In this study, EA supplementation increased the expression of occludin, Reg3β, BD2, MUC2, and BD1. These results were indicating that EA may improve colonic barrier functions via reducing gut permeability and increasing antimicrobial protein secretion.

IBD patients consistently suffered from chronic inflammation characterized by up-regulated proinflammatory cytokines such as IL1, IL6, IL17, TNFα, and INFγ that contribute to the establishment of tumor-associated immune microenvironment^14^. These proinflammatory cytokines are produced by CD4+ T helper (Th) lineage cells via IL12-Th1 and IL-23-Th17 pathways in IBD patients and mice with colitis^40–41^. Alterations in gut microbiota also have been demonstrated to be responsible for the regulation of Th1/Th17 immune response in IBD^2, 40^. Besides, sIgA up-regulated in the colon of IBD patients than that in healthy people^1^. Secretory IgA is generated by the combined function of plasma cells producing multimeric IgA and epithelial cells expressing pIgR, and it can protect the mucosal barrier against toxins and bacteria infections^42^. In this study, IL1, IL6, IL17, TNFα, and INFγ showed a significant reduction in EA treated groups. Conversely, sIgA was significantly up-regulated in EA treated groups. Further, IL12-Th1 and IL23-Th17 pathway-related cytokine gene expression were investigated. IL1β mRNA showed a significant reduction in EA treated groups. IL6, IL10, IL22, and sIgA mRNA were significantly up-regulated in the high EA treated group. However, relative TNFα mRNA expression showed a significant rise in EA treated groups. These results indicated that EA treatments may alleviate colonic inflammation via down-regulating both IL-12-Th1 and IL23-Th17 pathways and increase the sIgA secretion. ALT and AST are liver-specific enzymes released into serum following acute liver damage^29^. Both IBD patients and mice models have been demonstrated to hold high serum AST and ALT concentrations as a result of inflammatory liver injury^43^. In this research, both serum ALT and AST were significantly down-regulated in the high EA treated group in DSS-treated mice. These results indicated that EA may had a benefical impact on liver inflammation, further studies need to elucidate it.

To date, most EA signaling related researches were focused on bacterial utilization and CDP-EA pathway in PE synthesis^11, 17, 19–20^. Our previous study has demonstrated that ethanolamine could promote the proliferation of IPEC-J1 cells by regulating the mTOR signaling pathway and mitochondrial function^44^. Toll-like receptors (TLRs) have a vital role in mucosal immune responses to gut bacteria, and the TLR4 expression was always dramatically up-regulated in the intestines of IBD patients^21^. Myeloiddifferentiationfactor 88 holds its essential role in the regulation of innate gut immunity, and it is the direct downstream of TLRs and cytokine receptors^22^. Nutrient signals such as peptidoglycan and lipopolysaccharide can activate TLR4-MyD88 dependent or independent pathways to regulate the expression of antimicrobial proteins like the Reg3 protein family that ultimately reprogramme the gut microbiome in IBD^23–24^. Here, in the DSS-treated mice model, TLR4 protein was significantly up-regulated in EA treated groups, and MyD88 protein showed significantly up-regulated in the high EA treated group. However, MyD88 mRNA was up-regulated in the highEA group, while TLR4 only showed a significant increase in the low EA group. Crypt-derived colonic organoids can spontaneously generate crypt-villus like units and constituting by all types of intestinal cells. And so that colonic organoids hold the cellular and structural heterogeneity that highly coincident with the physiological nature of intestinal epithelium and much better than traditional cell lines in characterizing the immunology of intestinal barrier^37^. Hence, a DSS-induced colonic organoid inflammation model was established to verify these findings. Results showed that TLR4 protein expression tended to be more significant in the 500μM EA treated group. At the same time, TLR4 mRNA showed a significant increase in 500μM EA treated group.MyD88 mRNA expression was significantly up-regulated in both 100μM, and 500μM EA treated group. These results demonstrated that EA could exert impacts on the regulation of TLR4/MyD88 signals, and differences between mice model and organoid model may be caused by gut microbiota^3^, and EA may mediate host-microbiome cross-talk via TLR4/MyD88 dependent signaling in the inflamed gut.

In conclusion, this research demonstrated that EA may alleviate colonic inflammatory immunoreactions and microbiome dysbiosis via TLR4/MyD88 dependent signaling. EA treatment improved intestinal barrier functions by up-regulating the expression of occluding and antimicrobial protein. EA treatment alleviated colonic inflammatory response by down-regulating related cytokines (IL1, IL6, IL17, TNFα, and INFγ) and increasing sIgA secretion. Results from both DSS-treated mice and DSS-treated colonic organoids indicated that EA may directly target the TLR4/MyD88 dependent pathway to mediate host-microbial interactions in intestinal inflammation.

## Acknowledgments

Jian Zhou thanks professor Xiong, Wan, and Yin for their support and encouragement. Zhou, Xiong, and Yin thank Pan Huang for her technical support.

## Supplemental Materials

See the file “Supplemental materials. docx”

## References

1. Scaldaferri, F.; Gerardi, V.; Lopetuso, L. R.; Del Zompo, F.; Mangiola, F.; Boskoski, I.; Bruno, G.; Petito, V.; Laterza, L.; Cammarota, G.; Gaetani, E.; Sgambato, A.; Gasbarrini, A., Gut Microbial Flora, Prebiotics, and Probiotics in IBD: Their Current Usage and Utility. Biomed Research International 2013.

2. Wong, S. H.; Yu, J., Gut microbiota in colorectal cancer: mechanisms of action and clinical applications. Nat Rev Gastroenterol Hepatol 2019, 16 (11), 690–704.

3. Lavelle, A.; Sokol, H., Gut microbiota-derived metabolites as key actors in inflammatory bowel disease. Nat Rev Gastroenterol Hepatol 2020, 17(4), 223–237.

4. Cani, P. D.; Jordan, B. F., Gut microbiota-mediated inflammation in obesity: a link with gastrointestinal cancer. Nat Rev Gastroenterol Hepatol 2018, 15(11), 671–682.

5. Round, J. L.; Mazmanian, S. K., The gut microbiota shapes intestinal immune responses during health and disease. Nat Rev Immunol 2009, 9 (5), 313–23.

6. Honda, K.; Littman, D. R., The microbiome in infectious disease and inflammation. Annu Rev Immunol 2012, 30, 759–95.

7. Ren, W.; Chen, S.; Yin, J.; Duan, J.; Li, T.; Liu, G.; Feng, Z.; Tan, B.; Yin, Y.; Wu, G., Dietary arginine supplementation of mice alters the microbial population and activates intestinal innate immunity. J Nutr 2014, 144 (6), 988–95.

8. Gao, J.; Xu, K.; Liu, H.; Liu, G.; Bai, M.; Peng, C.; Li, T.; Yin, Y., Impact of the Gut Microbiota on Intestinal Immunity Mediated by Tryptophan Metabolism. Front Cell Infect Microbiol 2018, 8, 13.

9. Hamer, H. M.; Jonkers, D.; Venema, K.; Vanhoutvin, S.; Troost, F. J.; Brummer, R. J., Review article: the role of butyrate on colonic function. Aliment Pharmacol Ther 2008, 27 (2), 104–19.

10. Zhou, J.; Xiong, X.; Wang, K.; Zou, L.; Lv, D.; Yin, Y., Ethanolamine Metabolism in the Mammalian Gastrointestinal Tract: Mechanisms, Patterns, and Importance. Curr Mol Med 2017, 17 (2), 92–99.

11. Wu, Y.; Chen, K. S.; Xing, G. S.; Li, L. P.; Ma, B. C.; Hu, Z. J.; Duan, L. F.; Liu, X. G., Phospholipid remodeling is critical for stem cell pluripotency by facilitating mesenchymal-to-epithelial transition. Sci Adv 2019, 5 (11).

12. Patel, D.; Witt, S. N., Ethanolamine and Phosphatidylethanolamine: Partners in Health and Disease. Oxid Med Cell Longev 2017, 2017, 1–18.

13. Zhou, J.; Xiong, X.; Wang, K. X.; Zou, L. J.; Ji, P.; Yin, Y. L., Ethanolamine enhances intestinal functions by altering gut microbiome and mucosal anti-stress capacity in weaned rats. Br J Nutr 2018, 120 (3), 241–249.

14. Lin, Y. C.; Lin, Y. C.; Chen, C. J., Cancers Complicating Inflammatory Bowel Disease. N Engl J Med 2015, 373 (2), 194–5.

15. Brown, E. M.; Ke, X. B.; Hitchcock, D.; Jeanfavre, S.; Avila-Pacheco, J.; Nakata, T.; Arthur, T. D.; Fornelos, N.; Heim, C.; Franzosa, E. A.; Watson, N.; Huttenhower, C.; Haiser, H. J.; Dillow, G.; Graham, D. B.; Finlay, B. B.; Kostic, A. D.; Porter, J. A.; Vlamakis, H.; Clish, C. B.; Xavier, R. J., Bacteroides-Derived Sphingolipids Are Critical for Maintaining Intestinal Homeostasis and Symbiosis. Cell Host & Microbe 2019, 25(5), 668–+.

16. Thiennimitr, P.; Winter, S. E.; Winter, M. G.; Xavier, M. N.; Tolstikov, V.; Huseby, D. L.; Sterzenbach, T.; Tsolis, R. M.; Roth, J. R.; Baumler, A. J., Intestinal inflammation allows Salmonella to use ethanolamine to compete with the microbiota. Proc Natl Acad Sci U S A 2011, 108 (42), 17480–5.

17. Ormsby, M. J.; Logan, M.; Johnson, S. A.; McIntosh, A.; Fallata, G.; Papadopoulou, R.; Papachristou, E.; Hold, G. L.; Hansen, R.; Ijaz, U. Z.; Russell, R. K.; Gerasimidis, K.; Wall, D. M., Inflammation associated ethanolamine facilitates infection by Crohn’s disease-linked adherent-invasive Escherichia coli. EBioMedicine 2019, 43, 325–332.

18. Kaval, K. G.; Singh, K. V.; Cruz, M. R.; DebRoy, S.; Winkler, W. C.; Murray, B. E.; Garsin, D. A., Loss of Ethanolamine Utilization in Enterococcus faecalis Increases Gastrointestinal Tract Colonization. mBio 2018, 9 (3).

19. Anderson, C. J.; Clark, D. E.; Adli, M.; Kendall, M. M., Ethanolamine Signaling Promotes Salmonella Niche Recognition and Adaptation during Infection. PLoS Pathog 2015, 11 (11), e1005278.

20. Moore, T. C.; Escalante-Semerena, J. C., The EutQ and EutP proteins are novel acetate kinases involved in ethanolamine catabolism: physiological implications for the function of the ethanolamine metabolosome in Salmonella enterica. Mol Microbiol 2016, 99 (3), 497–511.

21. Cario, E.; Podolsky, D. K., Differential alteration in intestinal epithelial cell expression of toll-like receptor 3 (TLR3) and TLR4 in inflammatory bowel disease. Infect Immun 2000, 68 (12), 7010–7.

22. Koliaraki, V.; Chalkidi, N.; Henriques, A.; Tzaferis, C.; Polykratis, A.; Waisman, A.; Muller, W.; Hackam, D. J.; Pasparakis, M.; Kollias, G., Innate Sensing through Mesenchymal TLR4/MyD88 Signals Promotes Spontaneous Intestinal Tumorigenesis. Cell Reports 2019, 26 (3), 536–+.

23. Chu, H.; Mazmanian, S. K., Innate immune recognition of the microbiota promotes host-microbial symbiosis. Nat Immunol 2013, 14 (7), 668–75.

24. Chaniotou, Z.; Giannogonas, P.; Theoharis, S.; Teli, T.; Gay, J.; Savidge, T.; Koutmani, Y.; Brugni, J.; Kokkotou, E.; Pothoulakis, C.; Karalis, K. P., Corticotropin-releasing factor regulates TLR4 expression in the colon and protects mice from colitis. Gastroenterology 2010, 139 (6), 2083–92.

25. Zhou, J.; Xiong, X.; Yin, J.; Zou, L.; Wang, K.; Shao, Y.; Yin, Y., Dietary Lysozyme Alters Sow’s Gut Microbiota, Serum Immunity and Milk Metabolite Profile. Front Microbiol 2019, 10, 177.

26. Zhai, Z.; Zhang, F.; Cao, R.; Ni, X.; Xin, Z.; Deng, J.; Wu, G.; Ren, W.; Yin, Y.; Deng, B., Cecropin A Alleviates Inflammation Through Modulating the Gut Microbiota of C57BL/6 Mice With DSS-Induced IBD. Front Microbiol 2019, 10, 1595.

27. Miyoshi, H.; Stappenbeck, T. S., In vitro expansion and genetic modification of gastrointestinal stem cells in spheroid culture. Nat Protoc 2013, 8 (12), 2471–82.

28. Wen, Y. A.; Li, X.; Goretsky, T.; Weiss, H. L.; Barrett, T. A.; Gao, T., Loss of PHLPP protects against colitis by inhibiting intestinal epithelial cell apoptosis. Biochim Biophys Acta 2015, 1852 (10 Pt A), 2013–23.

29. Zou, L.; Xiong, X.; Liu, H.; Zhou, J.; Liu, Y.; Yin, Y., Effects of dietary lysozyme levels on growth performance, intestinal morphology, immunity response and microbiota community of growing pigs. J Sci Food Agric 2019, 99 (4), 1643–1650.

30. Xiong, X.; Zhou, J.; Liu, H.; Tang, Y.; Tan, B.; Yin, Y., Dietary lysozyme supplementation contributes to enhanced intestinal functions and gut microflora of piglets. Food Funct 2019, 10 (3), 1696–1706.

31. Asshauer, K. P.; Wemheuer, B.; Daniel, R.; Meinicke, P., Tax4Fun: predicting functional profiles from metagenomic 16S rRNA data. Bioinformatics 2015, 31 (17), 2882–4.

32. Manichanh, C.; Rigottier-Gois, L.; Bonnaud, E.; Gloux, K.; Pelletier, E.; Frangeul, L.; Nalin, R.; Jarrin, C.; Chardon, P.; Marteau, P.; Roca, J.; Dore, J., Reduced diversity of faecal microbiota in Crohn’s disease revealed by a metagenomic approach. Gut 2006, 55(2), 205–11.

33. O’Keefe, S. J. D.; Li, J. V.; Lahti, L.; Ou, J. H.; Carbonero, F.; Mohammed, K.; Posma, J. M.; Kinross, J.; Wahl, E.; Ruder, E.; Vipperla, K.; Naidoo, V.; Mtshali, L.; Tims, S.; Puylaert, P. G. B.; DeLany, J.; Krasinskas, A.; Benefiel, A. C.; Kaseb, H. O.; Newton, K.; Nicholson, J. K.; de Vos, W. M.; Gaskins, H. R.; Zoetendal, E. G., Fat, fibre and cancer risk in African Americans and rural Africans. Nature Communications 2015, 6.

34. Ni, J.; Wu, G. D.; Albenberg, L.; Tomov, V. T., Gut microbiota and IBD: causation or correlation? Nat Rev Gastroenterol Hepatol 2017, 14 (10), 573–584.

35. Fries, W.; Belvedere, A.; Vetrano, S., Sealing the Broken Barrier in IBD: Intestinal Permeability, Epithelial Cells and Junctions. Current Drug Targets 2013, 14 (12), 1460–1470.

36. Witten, J.; Samad, T.; Ribbeck, K., Selective permeability of mucus barriers. Curr Opin Biotech 2018, 52, 124–133.

37. Bar-Ephraim, Y. E.; Kretzschmar, K.; Clevers, H., Organoids in immunological research. Nat Rev Immunol 2020, 20 (5), 279–293.

38. Anhe, F. F.; Pilon, G.; Roy, D.; Desjardins, Y.; Levy, E.; Marette, A., Triggering Akkermansia with dietary polyphenols: A new weapon to combat the metabolic syndrome? Gut Microbes 2016, 7 (2), 146–53.

39. Everard, A.; Belzer, C.; Geurts, L.; Ouwerkerk, J. P.; Druart, C.; Bindels, L. B.; Guiot, Y.; Derrien, M.; Muccioli, G. G.; Delzenne, N. M.; de Vos, W. M.; Cani, P. D., Cross-talk between Akkermansia muciniphila and intestinal epithelium controls diet-induced obesity. P Natl Acad Sci USA 2013, 110 (22), 9066–9071.

40. Moschen, A. R.; Tilg, H.; Raine, T., IL-12, IL-23 and IL-17 in IBD: immunobiology and therapeutic targeting. Nat Rev Gastroenterol Hepatol 2019, 16 (3), 185–196.

41. Galvez, J., Role of Th17 Cells in the Pathogenesis of Human IBD. ISRN Inflamm 2014, 2014, 928461.

42. Pabst, O.; Slack, E., IgA and the intestinal microbiota: the importance of being specific. Mucosal Immunology 2020, 13 (1), 12–21.

43. Oliveira, G. R.; Teles, B. C. V.; Brasil, E. F.; Souza, M. H. L. P.; Furtado, L. E. T. A.; de Costro-Costa, C. M.; Rola, F. H.; Braga, L. L. B. C.; Gondim, F. D. A., Peripheral neuropathy and neurological disorders in an unselected Brazilian population-based cohort of IBD patients. Inflammatory Bowel Diseases 2008, 14 (3), 389–395.

44. Yang, H.; Xiong, X.; Li, T.; Yin, Y., Ethanolamine enhances the proliferation of intestinal epithelial cells via the mTOR signaling pathway and mitochondrial function. In Vitro Cell Dev Biol Anim 2016, 52 (5), 562–7.

